# E-cadherin tunes tissue mechanical behavior before and during morphogenetic tissue flows

**DOI:** 10.1101/2024.05.07.592778

**Authors:** Xun Wang, Christian M. Cupo, Sassan Ostvar, Andrew D. Countryman, Karen E. Kasza

## Abstract

Adhesion between epithelial cells enables the remarkable mechanical behavior of epithelial tissues during morphogenesis. However, it remains unclear how cell-cell adhesion influences mechanics in static as well as in dynamically flowing epithelial tissues. Here, we systematically modulate E-cadherin-mediated adhesion in the *Drosophila* embryo and study the effects on the mechanical behavior of the germband epithelium before and during dramatic tissue remodeling and flow associated with body axis elongation. Before axis elongation, we find that increasing E-cadherin levels produces tissue comprising more elongated cells and predicted to be more fluid-like, providing reduced resistance to tissue flow. During axis elongation, we find that the dominant effect of E-cadherin is tuning the speed at which cells proceed through rearrangement events, revealing potential roles for E-cadherin in generating friction between cells. Before and during axis elongation, E-cadherin levels influence patterns of actomyosin-dependent forces, supporting the notion that E-cadherin tunes tissue mechanics in part through effects on actomyosin. Taken together, these findings reveal dual—and sometimes opposing—roles for E-cadherin-mediated adhesion in controlling tissue structure and dynamics *in vivo* that result in unexpected relationships between adhesion and flow.

## Introduction

During embryonic development, epithelial tissue sheets play vital roles in physically supporting embryo structure, serving as boundaries and barriers, and dynamically remodeling to generate new tissue and organ structures through morphogenetic processes.^1–3^ These functions are associated with cohesive tissue structures comprising tightly packed cells that can remodel and flow at specific times and locations in the embryo. For example, the germband epithelium in the *Drosophila* embryo converges and extends through oriented cell rearrangements that double the length of the body axis in a rapid flow while maintaining tissue cohesion.^4–6^

The mechanical properties of epithelial tissues are central to their roles in development.^7–10^ Recent experimental studies have demonstrated that epithelial tissues can behave as solid-like materials that resist flow or fluid-like materials that accommodate flow.^11–15^ Theoretical and computational models have explored potential physical mechanisms underlying solid-fluid tissue behaviors in stable cellular packings.^16–19^ For example, in vertex models of epithelial tissues, the collective energy barrier for a group of cells to change shape and undergo a cell rearrangement depends on the preferred shapes of cells in the tissue, with high energy barriers associated with solid-like tissue behavior and low energy barriers with fluid-like behavior.^18–20^ Using an anisotropic vertex model framework to link stable cell packings to solid-fluid tissue behavior, our group and others showed that the *Drosophila* germband behaves as a solid-like tissue before axis elongation and becomes more fluid-like just prior to the onset of axis elongation.^13^ However, it remains unclear how solid-fluid tissue mechanical properties are tuned by the protein machineries in cells.

E-cadherin-based adhesion and actomyosin-generated contractile tension are thought to be the key machineries underlying tissue mechanics in many contexts.^7,10,21,22^ However, the molecular control of solid-fluid mechanical behavior in confluent tissues, such as the germband epithelium in *Drosophila*, remains poorly understood. Adhesive and contractile machineries are deployed at different levels and in distinct patterns across the developing *Drosophila* embryo. Prior to axis elongation, E-cadherin proteins form adhesive bonds between cells at adherens junctions and are essential for cohesion of the epithelium.^23,24^ E-cadherins couple to the actomyosin cytoskeleton to organize and transmit tension across tissues in many contexts.^25,26^ Experimental studies in other systems suggest that the balance between adhesion and tension determines the lengths of contacts between cells, with adhesion promoting long cell contacts, either directly through forming adhesive bonds or indirectly by down-regulating actomyosin, and contractile tension promoting short contacts.^21,27,28^ In this picture, the lengths of cell contacts in stable tissue packings, which are set by a balance between adhesion and tension, will control the shapes of cells. In the vertex model, cell shapes tune the energy barriers to cell rearrangement in stable tissue packings.^18,20^ In this framework, tissues comprising highly adhesive, elongated cells are predicted to have low rearrangement energy barriers and display fluid-like behavior. However, given the complex behavior and functions of cellular adhesive and contractile machineries, it is not clear if such relationships will hold.

During axis elongation, while the germband is rapidly rearranging and flowing, E-cadherin maintains a localization at adherens junctions. The forces driving tissue flow are thought to be provided in large part by myosin II, which is strongly enriched at junctions between anterior and posterior cell neighbors (AP edges) in a planar polarized fashion.^5,29–32^ As cells undergo rearrangements, adherens junctions at AP edges are disassembled and new contacts between dorsal and ventral cell neighbors (DV edges) are formed to allow cells to change their relative positions without opening gaps in the tissue.^4–6^ Experimental studies have revealed roles for E-cadherin in regulating actomyosin dynamics and junctional trafficking during cell rearrangement,^33–38^ indicating multiple potential effects of cell-cell adhesion on tissue dynamics in this context. Moreover, a theoretical apposed cortex adhesion model of epithelia that incorporate aspects of adhesion dynamics highlights that adhesion has the potential to provide frictional effects that slow rearrangements.^39^ It remains unclear how the level of adhesion will affect cell rearrangement rates in the tissue. Systematic experimental studies that link adhesion levels, cell packings, and tissue flow during dynamic morphogenetic processes will be required to answer these questions.

To investigate how E-cadherin-based cell-cell adhesion influences epithelial tissue mechanics *in vivo*, we systematically modulated E-cadherin levels in the primary epithelium of the *Drosophila* embryo and analyzed the effects on cell packings, tissue mechanics, and cell rearrangements in the germband before and during body axis elongation. We find effects of E-cadherin on germband epithelial tissue mechanics in stable tissue packings prior to axis elongation as well as in dynamically remodeling tissue during axis elongation, although these effects are relatively weak over a broad range of E-cadherin levels. Prior to axis elongation when the tissue is relatively static, increasing E-cadherin levels above wild type produces tissues comprising more elongated cells, consistent with the picture that adhesion contributes to preferred cell shapes, producing tissues predicted to have lower cell rearrangement energy barriers and be more-fluid-like. During axis elongation, increased E-cadherin levels are associated with slower steps in individual cell rearrangement events and influence the rate and extent of tissue elongation. In addition, we find that E-cadherin levels influence junctional myosin II and cell shape fluctuations, indicating that E-cadherin tunes tissue mechanics in part through effects on actomyosin. Taken together, our results demonstrate key roles for E-cadherin in tuning epithelial cell packings and cell rearrangement dynamics that result in unexpected effects of adhesion on tissue flow during body axis elongation.

## Results

### E-cadherin tunes the structure of cell packings in the germband epithelium prior to body axis elongation

To explore the role of cell-cell adhesion in epithelial tissue mechanics, we first investigated how the level of E-cadherin influences the structure and mechanics of the *Drosophila* germband epithelium prior to the onset of dramatic tissue movements associated with gastrulation and embryonic body axis elongation (Fig. 1A). To decrease E-cadherin levels, transgenic RNA interference (E-cad RNAi)^40^ and the Gal4/UAS system were used to partially inhibit E-cadherin expression in the embryo. To increase E-cadherin levels, transgenic E-cadherin was over-expressed (E-cad OE) using the Gal4/UAS system. To quantify the effects of these perturbations on E-cadherin levels at adherens junctions, which mediate cell-cell adhesion, we analyzed the intensity of E-cadherin in confocal images of fixed and stained embryos. Compared to E-cadherin levels in wild-type embryos, we found that E-cadherin levels visualized by immunofluorescence were decreased by half in RNAi and increased more than two-fold in OE embryos (Fig. 1C and 1G, flies raised at 23ºC).

**Figure 1.**
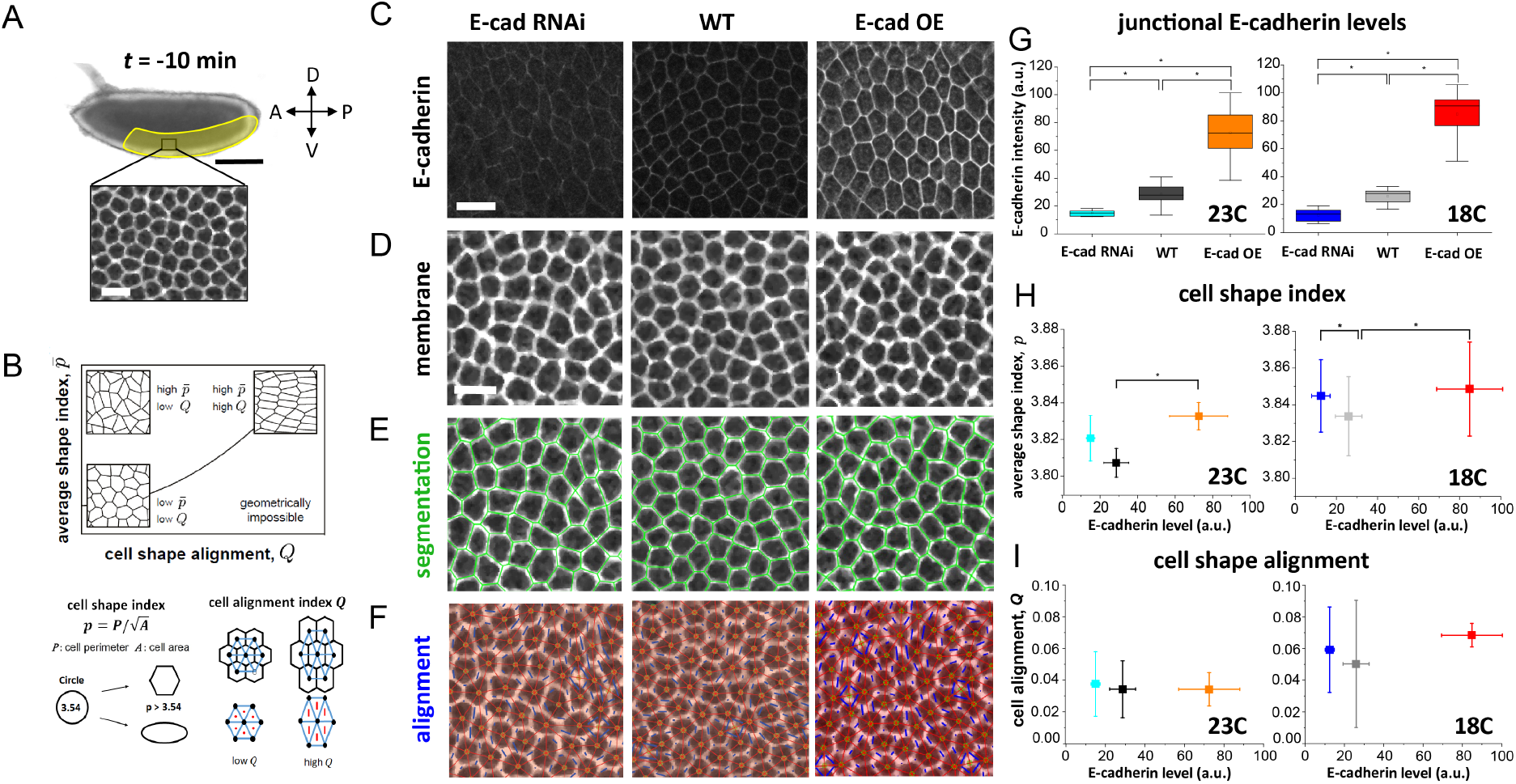
Cell shapes and packings in the germband prior to body axis elongation are influenced by E-cadherin levels. (A) Top: Image of *Drosophila* embryo 10 min before body axis elongation. Anterior, left. Ventral, down. Germband epithelial tissue is highlighted. Scale bar, 100 μm. Bottom: Germband epithelial cells with fluorescently labeled cell membranes. Scale bar, 10 μm. (B) Cell patterns in tissues can be characterized by the cell shape index *p* and cell shape alignment *Q*, figure modified with permission from Wang et al.^13^ The cell shape index, *p*, is calculated for each cell from the ratio of cell perimeter to square root of cell area, and the average value for cells in the tissue, 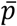, is calculated at each time point. Cell shape alignment *Q* is calculated from a triangulation of the tissue and quantifies how elongated and aligned cell shapes are across the tissue. High 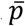 indicates elongated cell shapes, while *Q* is only high if elongated cell shapes are also aligned. (C) Confocal images of E-cadherin protein localization in the germband of wild-type (WT), E-cad RNAi, and E-cad OE embryos fixed and stained for E-cadherin in stage 6, prior to the onset of body axis elongation. Scale bar, 10 μm. (D-F) Confocal images of the germband ∼10 min prior to the onset of axis elongation in WT, E-cad RNAi, and E-cad OE embryos. Scale bars, 10 μm. (D) Cell membranes labeled with gap43:mCherry. (E) Overlaid polygon representation segmentation of epithelial cells (*green*). (F) Cell centers (*green dots*) are connected with each other by a triangular network (*red bonds*), which is then used to quantify cell stretching (*blue*) and alignment across the tissue. (G) Junctional E-cadherin levels in the germband of WT, E-cad RNAi, and E-cad OE embryos raised at 23ºC or 18ºC and fixed and stained for E-cadherin in stages 6 and 7. There were no statistically significant differences between stages 6 and 7, so embryos from these stages were grouped together here. n = 7-41 fixed embryos per group. (H) The average cell shape index 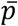 is tuned by E-cadherin levels. (I) Cell shape alignment *Q* is low and shows no dependence on E-cadherin levels. (H-I) n = 5-8 embryos per group. Error bars, standard deviation. P<0.05 denoted by a star.

To test how E-cadherin levels influence tissue structure, we analyzed cell shapes within the germband tissue of E-cad RNAi and OE embryos. The preferred shape of epithelial cells has been proposed to be determined by a balance between cell-cell adhesion, which promotes longer cell-cell contacts, and contractile tension, which promotes shorter contacts.^11,18,41,42^ In this picture, increased adhesion is predicted to produce longer cell-cell contacts and thus more elongated cell shapes if contractile forces are maintained constant.

To test how adhesion influences cell shapes, we analyzed segmented cell outlines in confocal images of embryos expressing a fluorescently tagged cell membrane marker using SEGGA image analysis software.^43^ We focused on a time point 10 min prior to the onset of germband extension, when the tissue is not being deformed by forces associated with gastrulation or body axis elongation. As a metric for whether cells have long or short contacts with neighbors, we quantified the average apical cell shape index, 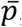, which is calculated from the cell perimeter *P* divided by the square root of the cell area 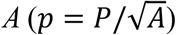 averaged over all cells in the region of interest (Fig. 1B). In wild-type embryos, cells were roughly isotropic and hexagonal in shape with short contacts with neighbors and compact shapes (Fig. 1D and 1E). The average cell shape index was 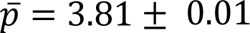 (Fig. 1H), consistent with prior reports.^13^ As an additional metric for quantifying cell packing structure in the tissue, we also quantified tissue anisotropy using the cell shape alignment index *Q*, calculated from the average cell elongation in the tissue^13,44,45^ (Fig. 1B) and found low alignment (Fig. 1F and 1I), consistent with prior reports indicating an isotropic tissue at this time point.^13^

First, we tested if increasing E-cadherin at cell-cell contacts influences cell shapes by quantifying 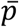 in the E-cad OE embryos. We found that E-cad OE cells were more elongated compared to wild type with 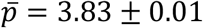 (Fig. 1D, E, and H, p-value of P = 0.002). This is consistent with reports from other systems in which cell-cell adhesion molecules tend to promote longer contacts between cells^21,46,47^ and with the simple picture that cell-cell adhesion promotes more elongated cell shapes, either directly through the energy associated with adhesive bonds or indirectly by down-regulating contractile tension. We found the value of the cell alignment index *Q* to be small in E-cad OE, similar to in wild type (Fig. 1F and 1I, P = 0.94), again consistent with an isotropic tissue.^13^

Next, we asked if the converse is true: does decreasing E-cadherin below wild-type levels produce less elongated cells? Unexpectedly, we found that 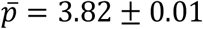 in E-cad RNAi embryos and was not decreased compared to in wild type (Fig. 1H, P = 0.07). Thus, we find an unexpected dependence of the cell shape index on E-cadherin levels—tissues with increased E-cadherin comprise cells that are more elongated than in wild-type embryos but tissues with decreased E-cadherin did not comprise cells that are less elongated.

To further explore how E-cadherin levels influence cell shapes, we utilized an inherent temperature dependence of the Gal4/UAS RNAi and over-expression methods to further increase or decrease E-cadherin levels by raising flies at 18ºC prior to transferring to room temperature for experiments. We found that the value of 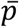 was larger in wild-type flies raised at 18ºC compared to 23ºC (Fig. 1G-I), suggesting a temperature dependence of cell packings that likely reflects effects of temperature on cytoskeletal dynamics and the overall rate of development during earlier embryonic stages. However, the overall dependence of 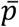 on E-cadherin levels persisted in flies raised at 18ºC or 23ºC, and the dependence was more clearly non-monotonic at 18ºC (Fig. 1G-I).

Taken together, these results demonstrate an unexpected dependence of cell shapes on E-cadherin levels. Our findings suggest that adhesion can promote longer cell-cell contacts and more elongated cell shapes, but that additional factors can also contribute to the cell packings in the germband.

### E-cadherin tunes the mechanical properties of the germband epithelium prior to body axis elongation

To test if E-cadherin levels influence tissue mechanical properties, we leveraged methods developed by our group and others to predict the solid-fluid mechanical behavior of epithelia from cell packings within the tissue.^13^ This link between cell shapes and tissue mechanics can be understood from vertex models of epithelia in which cell packings reflect the energy barriers to cell rearrangement, thereby revealing whether the model tissue behaves in a solid-like (finite energy barrier) or fluid-like (infinitesimal energy barrier) manner (Fig. 2A). If cells prefer to have short contacts with neighbors (small perimeters relative to areas), cells take on compact shapes, and it will take more energy to deform cells during rearrangements, associated with a higher energy barrier to rearrangement and more solid-like tissue behavior. In contrast, if cells prefer to have longer contacts with neighbors (large perimeters relative to areas), cells take on more elongated shapes, and it will take less energy to deform cells during rearrangements, associated with a lower energy barrier and fluid-like behavior. In this model, tissues comprising cells with more elongated shapes are predicted to be more fluid-like.

**Figure 2.**
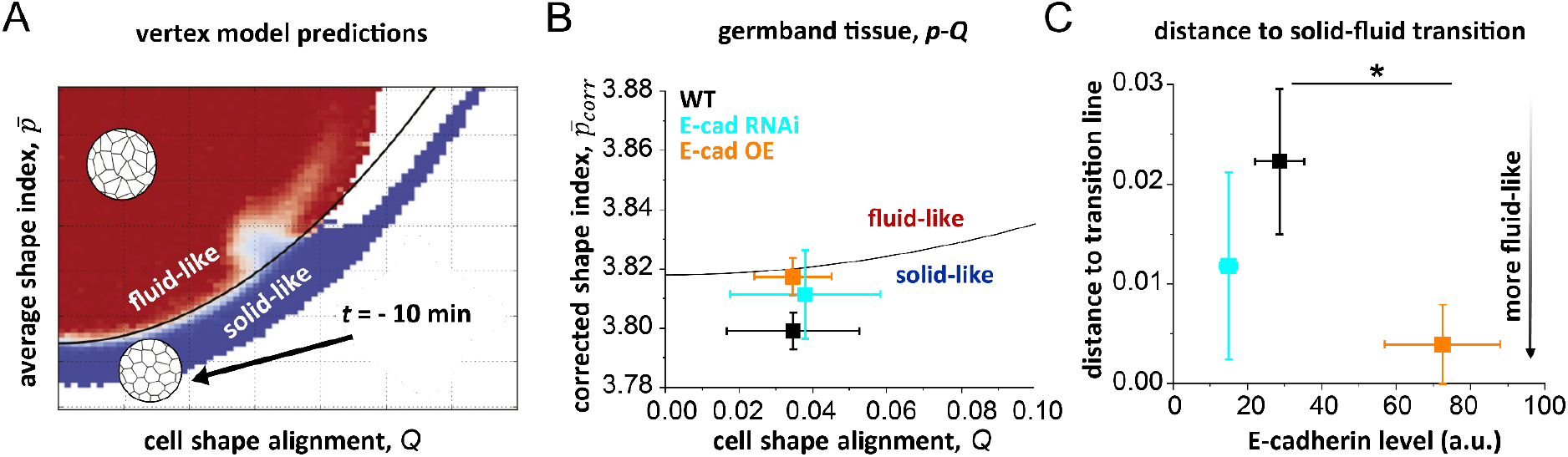
Germband tissue mechanical properties prior to axis elongation are influenced by E-cadherin levels. (A) Using a vertex model framework, solid-fluid tissue mechanical behavior is predicted from cell packings in the tissue characterized by the corrected cell shape index, 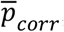, and cell shape alignment index, 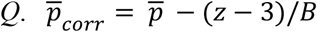, where z is the measured average vertex coordination number and *B* = 3.85 is a parameter previously determined from simulations.^48^ The predicted solid-fluid transition line (*black curve*) is given by 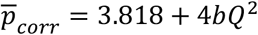, where *b* = 0.43.^13^ Figure adapted from Wang et al.^13^ (B) Predicted tissue mechanical behavior of the germband in WT (*black*), E-cad RNAi (*cyan*), and E-cad OE (*orange*) embryos based on cell packings 10 min before the onset of body axis elongation. In WT, the germband is predicted to be solid-like at this time point. The germband of embryos with increased (E-cad OE) or decreased (E-cad RNAi) E-cadherin levels are predicted to be solid-like but are positioned closer to the solid-fluid transition line, suggesting more fluid-like behavior. (C) Distance below the solid-fluid-like transition line is significantly reduced in the germband of E-cad OE (*orange*) compared to wild-type (*black*) embryos, n = 5-8 embryos per group.

For tissues like the germband that display some anisotropy, the average cell shape index 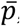, cell shape alignment index *Q*, and vertex coordination number *z*, are sufficient to classify a given packing as either solid-like or fluid-like in the vertex model framework. The predicted solid-fluid transition line (Fig. 2A, solid line) is given by 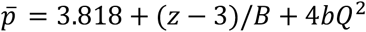, or alternatively, 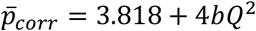, where *B* = 3.85, *b* = 0.43 and 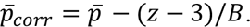.^13,48^ If the tissue sits above the line, it is predicted to behave as a fluid-like material, while the tissue is predicted to be solid-like if it is below the line (Fig. 2A). Comparing the cell packing parameters we measured in wild-type embryos to the predictions of the anisotropic vertex model,^13^ suggests that at this time point the tissue is positioned in the solid-like region of the *p-Q* parameter space, just below the solid-fluid transition line (Fig. 2B, *black*), consistent with prior work and with the observation that there are very few cell rearrangements during this time period.^13^

Next, we asked how E-cadherin levels influence solid-fluid tissue mechanical properties within the vertex model framework. Increased E-cadherin levels in E-cad OE embryos are associated with cell packings that are closer to the predicted solid-fluid transition line (Fig. 2B and 2C, *orange*, P = 0.046), suggesting more fluid-like mechanical behavior, consistent with the notion that adhesion might influence the collective energy barriers to cell rearrangement in the vertex model. Similarly, decreased E-cadherin levels in E-cad RNAi embryos are associated with a shift towards the transition line and more fluid-like behavior (Fig. 2B and 2C, *cyan*, P=0.37), although the change did not rise to the level of statistical significance. Therefore, higher or lower E-cadherin levels produce tissues predicted to be more fluid-like than in wild-type embryos.

### E-cadherin influences actomyosin localization patterns prior to body axis elongation

To investigate the mechanisms underlying the relationships between E-cadherin, cell shapes, and tissue mechanics, we explored if E-cadherin levels influence actomyosin in this context. Prior studies have demonstrated that not only does actomyosin contractility influence E-cadherin, but that E-cadherin can play a role in regulating actomyosin activity.^34,35,38,47,49–53^ Because the adhesive and contractile machineries are tightly coupled in many contexts, we reasoned that the observed relationships between E-cadherin levels and cell shapes might be explained if E-cadherin levels influence contractile tension in addition to directly mediating adhesion at cell-cell contacts.

To quantify changes in actomyosin levels, we imaged fluorescently tagged myosin II regulatory light chain in the wild-type, E-cad RNAi, and E-cad OE embryos. We found that increased E-cadherin in E-cad OE embryos was associated with a significant 60% decrease in myosin II levels at cell junctions compared to in wild-type embryos (Fig. 3A,B, P = 3.9 × 10^−4^), consistent with a previous report.^35^ By comparison, the decreased E-cadherin levels in E-cad RNAi embryos were associated with a 20% decrease in junctional myosin II compared to in wild type, although this change did not rise to the level of statistical significance (Fig. 3A,B, P = 0.27). Decreased junctional myosin II at cell-cell contacts would be predicted to lead to decreased contractile tension around the perimeter of cells and more elongated cell shapes. Consistent with this, the average cell shape index decreases with junctional myosin II levels in wild-type, E-cad RNAi, and E-cad OE embryos (Fig. 3C). Taken together, these results indicate that E-cadherin levels influence actomyosin localization at cell-cell contacts. This coupling between the adhesive and contractile machineries may help explain the observed relationships between E-cadherin, cell shapes, and tissue mechanics, suggesting a role for actomyosin in mediating the effects of E-cadherin on tissue mechanical behavior in this context.

**Figure 3.**
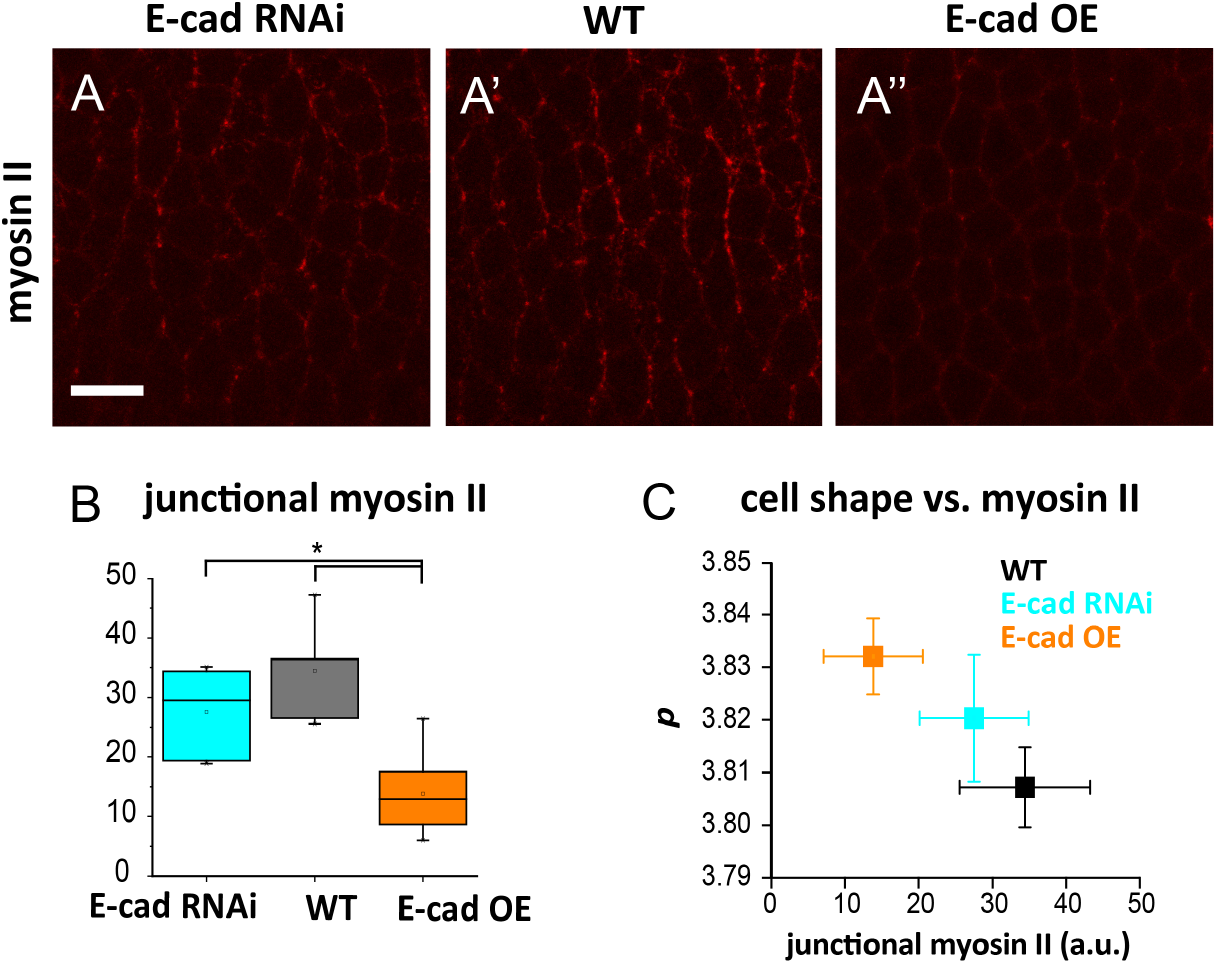
E-cadherin levels influence myosin II localization prior to body axis elongation. (A) Live confocal images of mCherry-tagged myosin II in the germband prior to the onset of body axis elongation. Scale bar, 10 μm. (B) Junctional myosin II intensities (a.u.) in WT, E-cad RNAi, and E-cad OE embryos. Junctional myosin II levels are decreased in E-cad OE compared to WT embryos, n = 11-18 embryos per group. (C) Average cell shape index vs. junctional myosin II levels in WT, E-cad RNAi, and E-cad OE embryos.

### Effects of E-cadherin levels on cell shapes during body axis elongation

We wondered how E-cadherin levels influence tissue structure and dynamics in the next phase of development, when the germband rapidly elongates and remodels through cell rearrangements during body axis elongation (Fig. 4A). In wild-type embryos, cells become more elongated during axis elongation, suggesting more fluid-like behavior.^13^

**Figure 4.**
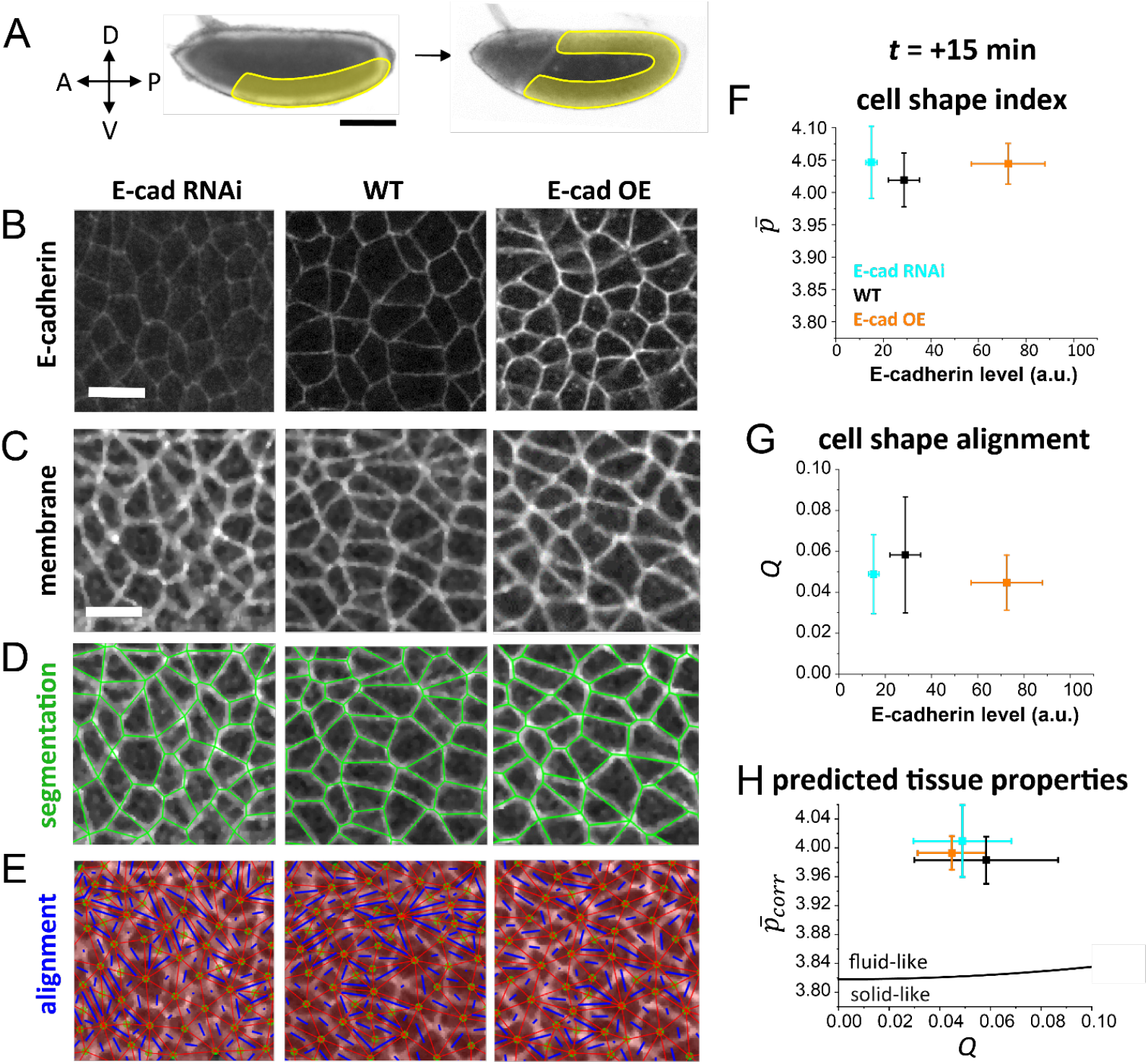
Effects of E-cadherin on cell packings during axis elongation. (A) Images of the *Drosophila* embryo before and after body axis elongation. The germband epithelium (*yellow*) narrows along the DV axis and elongates along the AP axis in a rapid tissue flow. Scale bar, 100 μm. Unless otherwise noted, *t* = 0 is defined as the time when body axis elongation begins. (B) Confocal images of germband cells in WT, E-cad RNAi, and E-cad OE embryos fixed and stained for E-cadherin at stage 7 during axis elongation. Scale bar, 10 μm. (C-E) Confocal images of live embryos at *t* = +15 min after the onset of body axis elongation, showing germband cells with cell membranes labeled with gap43:mCherry. Scale bar, 10 μm. (D) Overlaid segmentation of cells (*green*). (E) Overlaid triangulation for quantifying cell shape alignment. (F-H) Cell packings and predicted tissue mechanical behavior 15 min after the onset of body axis elongation in WT, E-cad RNAi and E-cad OE embryos. E-cadherin levels reflect those measured in fixed embryos and quantified in Fig. 1G. (F) The average cell shape index 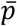. (G) The cell alignment index *Q*. (H) Tissue solid-fluid mechanical behavior based on vertex model predictions. In WT, E-cad RNAi, and E-cad OE embryos, the germband tissue is predicted to behave in a fluid-like manner at this time in development, n = 5-8 embryos per group.

We found that cell patterns in embryos in E-cad RNAi and E-cad OE embryos display a similar overall trend compared to those in wild type (Fig. 4B-G). Notably, the value of the cell shape index 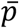 displayed qualitatively similar E-cadherin-dependent differences as we observed prior to axis elongation, but the observed differences were not statistically significant (Fig. 4F, P = 0.47). We did not observe defects in posterior midgut invagination in any of the embryo groups, which suggests that any observed cell shape changes in E-cad OE and E-cad RNAi embryos likely arise from cell autonomous effects in the germband instead of non-cell autonomous effects associated with posterior midgut invagination.^54,55^ The predicted mechanical behavior in E-cad RNAi, E-cad OE, and wild-type embryos were all fluid-like (Fig. 4H, P = 0.35). Taken together, these results indicate that E-cadherin levels may influence cell packings and predicted tissue mechanics in the germband while the tissue is actively flowing, although the effects appear less significant than when the tissue is largely static prior to axis elongation.

### E-cadherin levels influence the speed of cell rearrangement events during axis elongation

In addition to effects on cell shape and cell rearrangement energy barriers, E-cadherin-based adhesions can also influence the dynamics of cell rearrangements by giving rise to friction between cells or altering junctional disassembly/assembly processes. To explore if E-cadherin levels tune the speed of cell rearrangement events during axis elongation, we measured the timescales of distinct cell rearrangement steps in wild-type, E-cad RNAi, and E-cad OE embryos.

In wild-type embryos, a typical cell rearrangement is initiated by the contraction of a cell-cell contact between anterior and posterior cell neighbors (AP edge), giving rise to a higher-order vertex at which four or more cells meet^4–6^ (Fig. 5A). In wild-type embryos, the average rate of contraction was 1.4±0.4 μm/min (Fig. 5C), similar to previous reports.^56^ In the apposed cortex adhesion model,^39^ E-cadherin contributes to friction between cells and this friction can impede the rate of cell-cell contact contraction in some contexts, so we would predict decreased E-cadherin levels to be associated with faster contraction. In another picture, if E-cadherin is involved in cell junction disassembly/trafficking and if these processes mediate contraction,^34,35,37^ then we would predict decreased E-cadherin levels to be associated with slower contraction. We find that the contraction rate is reduced by 14% in E-cad RNAi embryos (1.2 ± 0.3 μm/min, P = 6.3 × 10^−4^) and by 10% in E-cad OE embryos (1.3 ± 0.4 μm/min, P = 0.02) compared to in wild type (Fig 5C). Consistent with these trends, the fraction of “vertical” AP cell contacts that do not contract to a vertex is increased in both E-cad RNAi and OE embryos (Fig 5B). Thus, increased or decreased E-cadherin levels at cell contacts is associated with slower contact contraction, suggesting multiple potential roles for E-cadherin in cell-cell contact contraction that are not consistent with either model alone.

**Figure 5.**
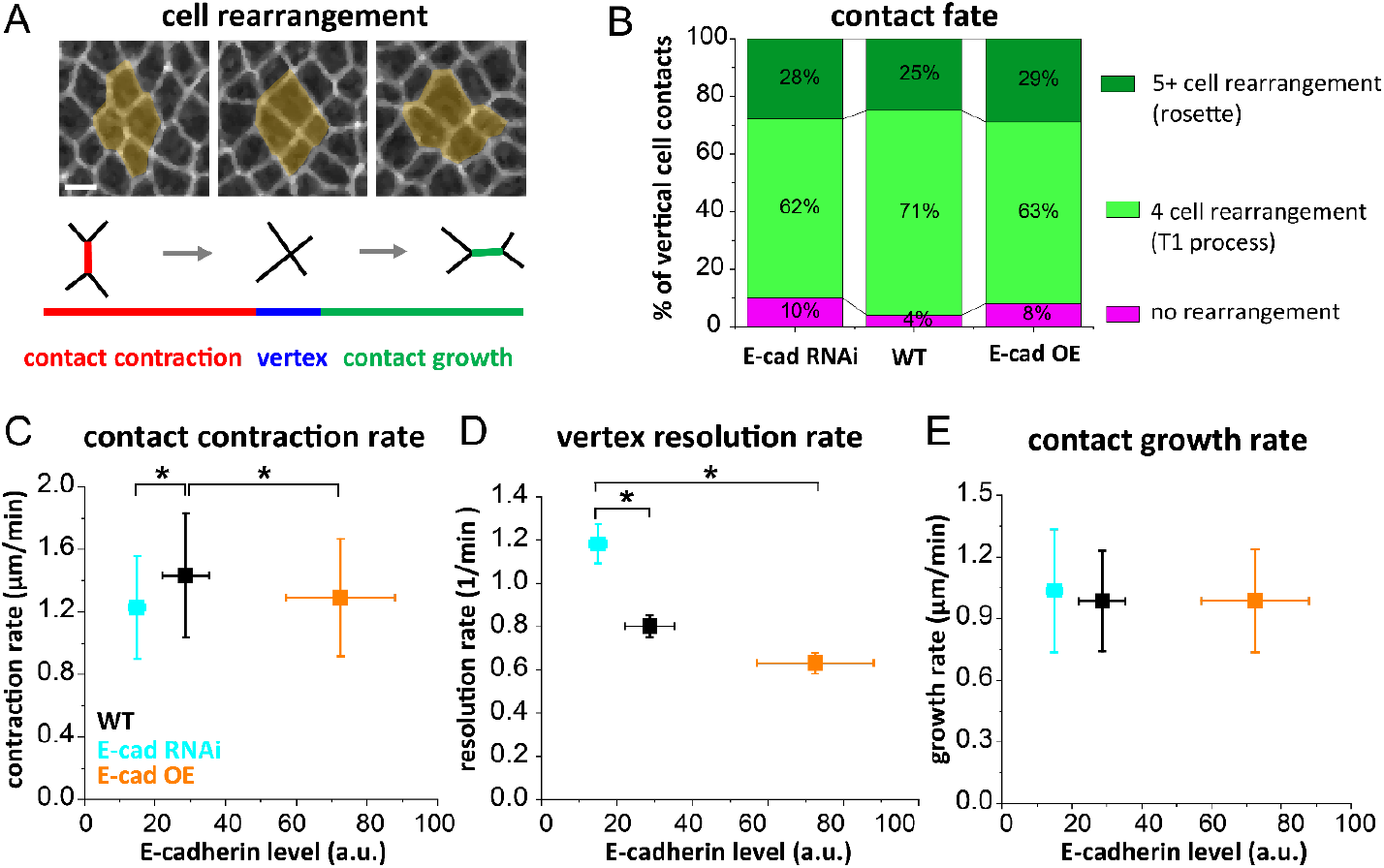
E-cadherin levels influence the speed of cell rearrangement events during body axis elongation. (A) Top: Still images from a confocal time-lapse movie of germband cells with fluorescently labeled cell membranes showing a cell rearrangement event. Scale bar, 5 μm. Bottom: Schematic of the steps in a typical cell rearrangement event. In the first step, rearrangement is initiated by contraction of a cell-cell contact, bringing four or more cells together at a high-order vertex. In the second step, the vertex exists for a short period of time before it resolves by formation of a short new contact between cells; the time that cells stay in the high-order vertex configuration is called the vertex lifetime. In the third step, the new contact grows to form a full new edge. (B) Fraction of “vertical” AP edges that participate in no rearrangement, a T1 rearrangement process involving 4 cells, or a rosette rearrangement involving 5 or more cells in WT, E-cad RNAi, and E-cad OE embryos, n = 100 edges per group. (C) Contact contraction rate is reduced in E-cad RNAi and E-cad OE compared to WT, n = 5-8 embryos per group. (D) The vertex resolution rate (1 / vertex lifetime) is increased in E-cad RNAi compared to WT, n = 484-759 cell rearrangements per group. Error bar, standard error. (E) The new contact growth rate is not significantly different between groups, n = 5-8 embryos per group.

Once a cell contact has contracted to a high-order vertex in wild-type embryos, this vertex then resolves to form a new contact between dorsal and ventral cell neighbors (DV edge), contributing to tissue elongation along the AP axis (Fig. 5A).^4–6,57^ The vertex resolution rate (1 / vertex lifetime) in wild-type embryos was 0.8 ± 1.4 min^−1^ (Fig. 5D). We found that reduced E-cadherin in E-cad RNAi embryos was associated with a 48% increase in vertex resolution rate (1.18 ± 2.04 min^−1^, P = 6 × 10^−5^), while increased E-cadherin in E-cad OE embryos was associated with a 21% decrease in this rate (0.63 ± 1.08 min^-1^, P = 0.14) compared to wild type (Fig. 5D). These results suggest that the levels of E-cadherin at cell-cell contacts tune vertex lifetime, with increased E-cadherin associated with slower vertex resolution, perhaps by influencing the timescale for disassembling and removing E-cadherin molecules from cells that were initially in contact or by influencing cellular machineries involved in forming the new cell-cell contact.

New cell contact growth involves the formation of new adhesive bonds between cells that were not previously in contact. This contact growth can occur passively in some models,^58^ while experimental studies have implicated actomyosin contractility in this process in some contexts.^57^ We did not find significant differences in the contact growth rates between wild-type, E-cad RNAi, and E-cad OE embryos (Fig. 5E).

Thus, we find that E-cadherin levels affect the speed of two distinct steps in cell rearrangements: contact contraction and vertex resolution. These effects have the potential to tune the rates of tissue remodeling and flow. The two contributions will both produce slower rearrangements in E-cad OE but have opposing effects in the E-cad RNAi case.

### E-cadherin influences actomyosin patterns and cell area fluctuations during axis elongation

We next explored if E-cadherin levels might also influence cell rearrangements through effects on actomyosin and contractile activity during axis elongation. We reasoned that E-cadherin might influence actomyosin during axis elongation in a manner that tunes the timescales of cell rearrangement events.

We find that the overall levels of myosin at junctions and the medial-apical domain were not significantly different in E-cad RNAi and E-cad OE compared to wild-type embryos during axis elongation (Fig. 6A-C), suggesting that the effects on myosin reported in other studies might be explained by the strength of the distinct perturbations.^35^ We did find that the planar polarized enrichment of myosin at “vertical” AP edges compared to “horizontal” DV edges was somewhat enhanced in E-cad OE compared to wild-type embryos (Fig. 6D, P = 0.04), potentially influencing forces driving cell rearrangement.

**Figure 6.**
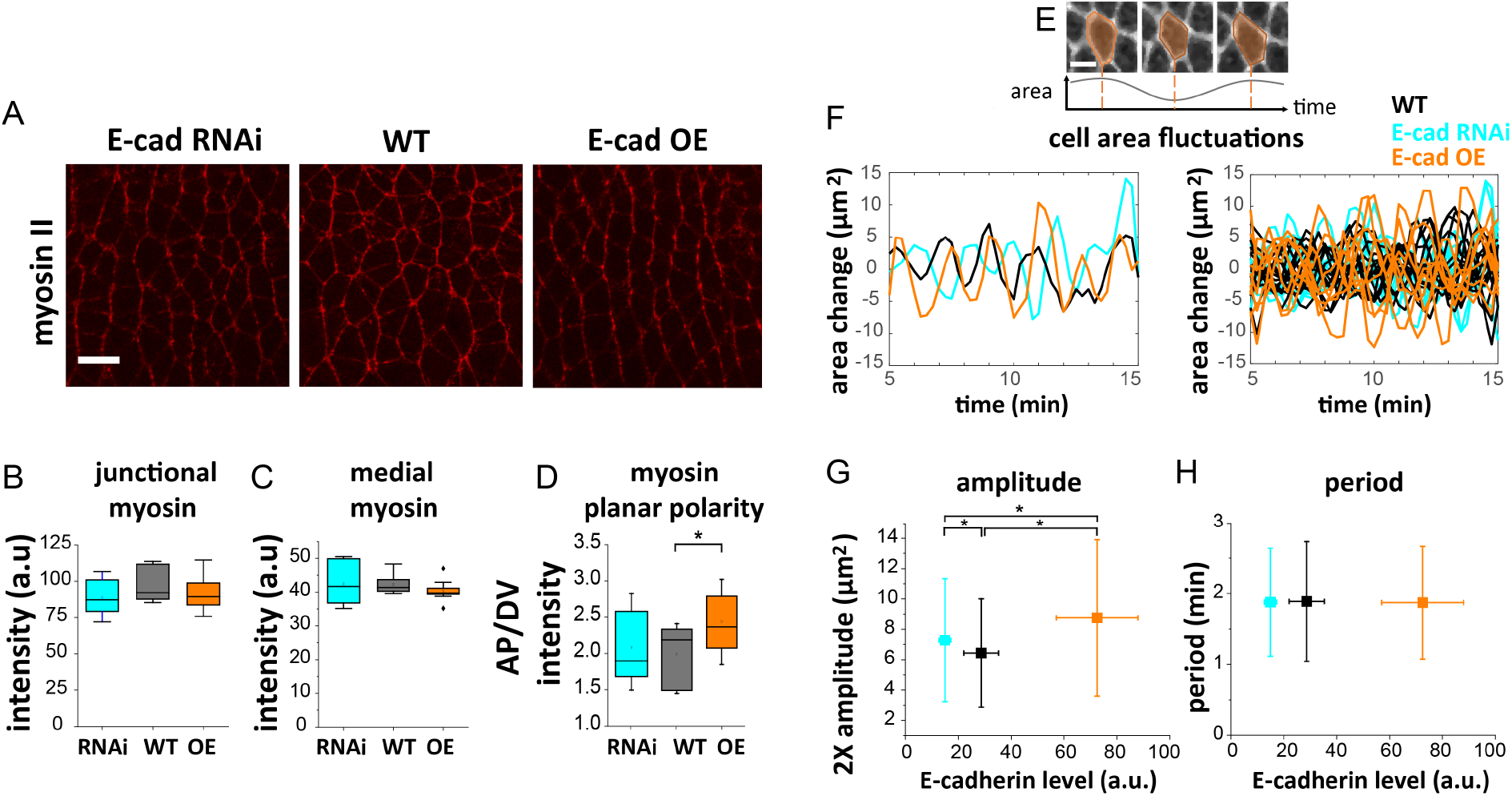
E-cadherin levels influence actomyosin patterns and cell area fluctuations during axis elongation. (A) Still images from timelapse confocal images of mCherry-tagged myosin II during body axis elongation in WT, E-cad RNAi, and E-cad OE embryos. Scale bar, 10 μm. (B) Junctional myosin II intensities. (C) Medial-apical myosin II intensities. (D) Planar polarity quantified from the ratio of myosin intensities at AP relative to DV cell junctions. n = 11-18 embryos per group. (E) Still images from a timelapse confocal movie illustrating apical cell area fluctuations in a cell of a WT embryo. Scale bar, 5 μm. (F) Left: Examples of cell area change (the difference in current cell area compared to the value 60 s prior) fluctuations over time in WT, E-cad RNAi, and E-cad OE embryos. Right: Cell area change fluctuations for all analyzed cells. Curves smoothed with a Savitzky-Golay filter. (G) The amplitude of cell area fluctuation is increased in E-cad RNAi and E-cad OE compared to in WT embryos, n = 482-516 area fluctuation cycles per group. (H) The period of cell area fluctuation is not significantly different between groups, n = 283-367 area fluctuation cycles per group.

Actomyosin contractility can generate pulsed contractions that drive apical cell area fluctuations,^33,57,60,61^ which have been proposed to help cells overcome energy barriers to cell rearrangement.^22,62,63^ We wondered if the effects of E-cadherin on actomyosin patterns might influence active fluctuations in this context. To explore this possibility, we measured apical cell area fluctuations in wild-type, E-cad RNAi, and E-cad OE embryos during axis elongation (Fig. 6E-H). In wild-type embryos, we observed apical cell area fluctuations with a mean period of 1.89 ± 0.85 min and amplitude of 6.4 ± 3.6 μm^2^ (Fig. 6G, H), consistent with prior studies.^61,64^

We did not find significant changes in the period in E-cad RNAi or OE embryos (Fig. 6H). However, the amplitude of fluctuations was increased in E-cad OE embryos (P = 4.8 × 10^−9^) and E-cad RNAi embryos (P = 0.01) compared to wild type (Fig. 6G). As we did not find significant differences in the levels of medial myosin between wild-type, E-cad RNAi, and E-cad OE embryos (Fig. 6C), this suggests that the differences in amplitude may be associated with other aspects of actomyosin dynamics or tissue mechanics that are altered by E-cadherin. Taken together, these results reveal a role for E-cadherin in regulating the actomyosin and pulsatile contractile activity of cells during axis elongation, which has the potential to influence the dynamics of cell rearrangement events and tissue-level mechanics.

### E-cadherin levels influence embryonic development

Finally, we wondered how E-cadherin levels ultimately influence the tissue-level flow that elongates the anterior-posterior body axis. In wild-type embryos, the germband in 98% of embryos fully elongates in 30 min, and 95% of embryos hatch to larvae (Fig. 7A-C). In E-cad RNAi embryos, 37% of embryos fail during cellularization and do not form gastrula (Fig. 7A). Of those embryos that initiate axis elongation, 92% fully elongate (Fig. 7A). Of the embryos that fully elongate, 71% go on to hatch (Fig. 7A and 7B). These defects associated with E-cad RNAi were rescued by expressing a transgenic E-cadherin, indicating that E-cadherin was the dominant protein affected by the RNAi. When E-cadherin levels are further decreased by raising the flies at a lower temperature, the defects in cellularization and axis elongation are significantly increased, so that only 28% fully elongate and 15% hatch (Fig. 7A and 7B).

**Figure 7.**
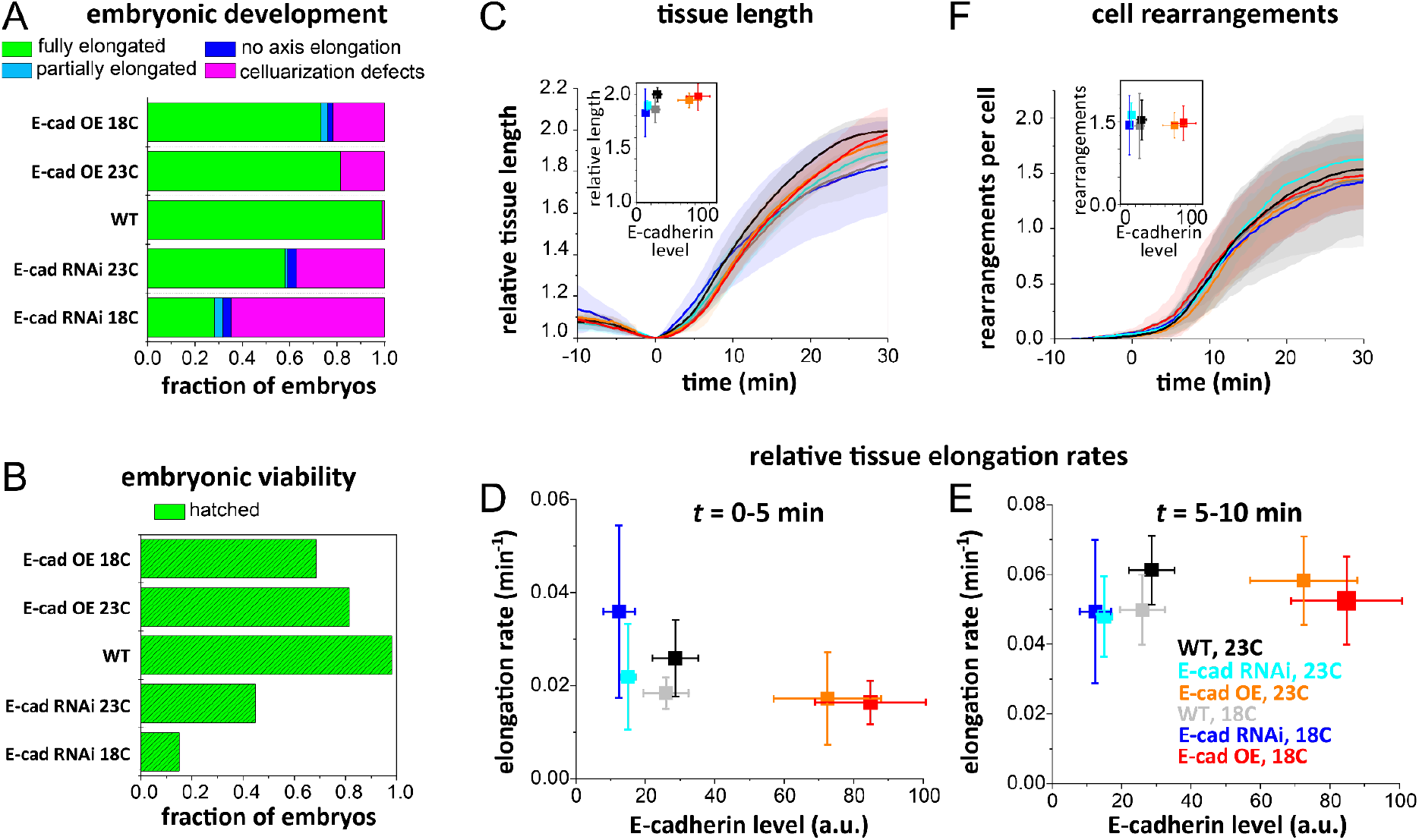
E-cadherin levels influence embryonic development. (A,B) Embryonic development defects and viability in WT, E-cad RNAi, and E-cad OE embryos. (A) Fraction of embryos that fail to cellularize, fail to initiate body axis elongation, only partially elongate, or fully elongate. (B) Fraction of embryos that hatch to larvae. n = 100 embryos per group. (C) In embryos that reach this stage of development and initiate body axis elongation, the germband epithelium rapidly doubles in length in about 30 min. Inset, the extent of tissue elongation at *t* = 30 min, n = 5-8 embryos per group. (D) Mean tissue elongation rates during the first 5 min of axis elongation decrease with increasing E-cadherin levels. (E) Mean tissue elongation rates from 5-10 min do not strongly vary with E-cadherin levels. (F) The cumulative number of cell rearrangements in the germband is not significantly different between WT, E-cad RNAi, and E-cad OE embryos.

Similarly, in E-cad OE embryos, 19% of embryos fail during cellularization stages and do not form gastrula (Fig. 7A), but of those that initiate axis elongation, 100% fully elongate (Fig. 7A). Of embryos that fully elongate, 100% go on to hatch (Fig. 7B). When E-cadherin levels are further increased by raising the flies at a lower temperature, the defects in cellularization and axis elongation are increased, so that 73% fully elongate and 69% hatch (Fig. 7A and 7B).

Decreasing E-cadherin levels by 50% produces a higher percentage of embryos with defects in tissue elongation (72%) and hatching (83%) than increasing E-cadherin levels more than 2-fold (25% and 32%, respectively) (Fig. 7A and 7B). Taken together, these results indicate that decreasing or increasing E-cadherin levels compared to wild type results in developmental defects, suggesting that there is an optimal range of E-cadherin levels for development but that this range is relatively large.

In embryos in which body axis elongation initiates, we find that the initial tissue elongation rate at the onset of axis elongation is linked with E-cadherin levels, with higher E-cadherin levels associated with slower tissue elongation during this time (Fig. 7C and D, *t* = 0-5 min), potentially reflecting the effects of E-cadherin on frictional contributions or actomyosin-dependent forces in cell rearrangements. Later in the process, tissue elongation rates do not appear to vary strongly with E-cadherin levels (Fig. 7E, *t* = 5-10 min), suggesting that the differences we observe in cell rearrangement speeds do not significantly affect the numbers of rearrangements initiated or the contribution to tissue flow (Fig. 7F). These effects combined lead to ∼5% and ∼3% decreases in final tissue length compared to wild type in E-cad RNAi and OE, respectively (Fig. 7C, *t* = 30 min, P = 0.03).

Taken together, these results reflect that axis elongation is influenced by but relatively tolerant to absolute changes in E-cadherin levels over a broad range, and that embryonic development overall is disrupted by changes in E-cadherin levels outside this range.

## Discussion

To investigate how E-cadherin-based cell-cell adhesion influences epithelial tissue mechanics *in vivo*, we modulated E-cadherin levels over a ∼4-fold range in the *Drosophila* primary epithelium and analyzed the effects on cell packings, tissue mechanics, and cell rearrangements in the germband before and during body axis elongation. We find that prior to axis elongation when the tissue is relatively static, increasing E-cadherin levels above wild-type levels produces tissue comprising more elongated cells and that is predicted to be more fluid-like, consistent with the picture that adhesion contributes to cell shapes in stable packings. In contrast, decreasing E-adherin levels below wild-type levels does not produce tissue comprising less elongated cells, suggesting that additional factors contribute to cell shape in this context. During axis elongation, increasing E-cadherin levels is associated with slower edge contraction and vertex resolution in cell rearrangement events, consistent with a role for E-cadherin in rearrangement dynamics, via direct frictional effects and/or indirect effects associated with regulation of actomyosin. These effects on rearrangement speed tune overall rates of tissue flow, but relatively weakly, suggesting a wide range of E-cadherin levels that support axis elongation. In addition, we find that E-cadherin levels influence junctional myosin II and active cell shape fluctuations, indicating that E-cadherin tunes tissue mechanics in part through effects on actomyosin and contractile tension in the tissue. Taken together, these results reveal dual roles for E-cadherin in tuning tissue mechanics—in lowering the cell shape-dependent energy barriers to cell rearrangement and in contributing to friction between and/or forces on cells as they move during rearrangements. These dual roles can have opposing effects during tissue flows, leading to unexpected relationships between adhesion and flow during axis elongation, which are not captured by existing vertex models that focus on the mechanics of stable packings of cells or apposed cortex adhesion models that include dynamic features of cell-cell adhesion.

Increasing E-cadherin levels ∼2-fold above wild-type levels in E-cad OE embryos produces an epithelium prior to axis elongation that comprises cells with longer contacts with neighbors (higher cell shape index *p*). This is consistent with a role for E-cadherin in increasing contact lengths between cells, directly through adhesive binding energies and/or indirectly by locally downregulating contractile tension in the cortex.^21,28,47^ Our observation of reduced junctional myosin II in E-cad OE tissue supports the notion that contractile tension is reduced by junctional E-cadherin, consistent with prior studies.^35^ Notably, decreasing E-cadherin levels to half of wild type in E-cad RNAi embryos *does not* produce an epithelium comprising cells with shorter contacts with neighbors. Rather we see a subtle increase in *p* in this context as well, indicating that the relationship between E-cadherin levels and cell shape index may not be monotonic. There are several possible explanations for this observation. First, this could be explained if the reduction in junctional E-cadherin levels in E-cad RNAi embryos does not result in a corresponding decrease in adhesion strength. Second, this could be explained if reduced E-cadherin levels influence actomyosin contractile tensions in E-cad RNAi cells. Indeed, we observe a subtle decrease in junctional myosin II in E-cad RNAi cells that is consistent with the observed changes in cell shapes, again supporting a role for E-cadherin in regulating myosin-dependent contractile tensions. Alternatively, there may be a constraint on how much *p* can be reduced in a solid-like tissue, given that the minimum value is *p* = 3.72, associated with a perfect hexagonal lattice.^18^

Consistent with the possibility of other constraints affecting cell packings, we find that the temperature at which flies are raised during embryogenesis influences cell packings in wild-type embryos. One potential explanation for this is that temperature affects cytoskeletal and developmental dynamics during the process of cellularization, which forms the primary epithelium and produces the initial cell packings in the tissue. This suggests that the history of the tissue, along with current adhesion and contractile tension levels, sets initial cell packings. During axis elongation when the tissue is rapidly remodeling and flowing, cell packings show similar trends as prior to axis elongation but are not statistically distinguishable between wild-type, E-cad RNAi, and E-cad OE embryos. This suggests that cell packings *during* body axis elongation may not be as strongly affected by absolute E-cadherin levels or that cell shapes during body axis elongation are more reflective of additional factors, such as the dynamic responses associated with tissue shape changes driven by internal forces and external forces. Future efforts to more accurately model dynamic features of convergent extension will help to distinguish between these possibilities.

Based on the changes in initial germband cell packings in E-cad OE embryos, the vertex model framework predicts that these tissues will have reduced energy barriers to cell rearrangement and be more fluid-like (but still generally solid-like) compared to wild-type tissue. Based on this, one might predict that the more elongated cells in E-cad OE reflect more fluid-like initial tissue properties that would translate into more rapid cell rearrangements and tissue flow during body axis elongation. However, we observed somewhat *slower* cell rearrangements and tissue elongation in E-cad OE compared to wild-type embryos. In principle, this could reflect stronger mechanical resistance to flow (higher tissue viscosity) or lower driving forces. Because we do not observe significant reductions in myosin levels and in fact observed increased myosin planar polarity and cell area fluctuations during body axis elongation, this suggests increased resistance to flow, inconsistent with our prediction based on initial cell packings just prior to axis elongation. This points toward potential additional roles for E-cadherin in tissue flow during dynamic tissue remodeling.

In addition to influencing cell packings and energy barriers to cell rearrangements in stable cell packings, E-cadherin-based cell-cell adhesion has the potential to contribute to frictional effects in dynamically remodeling tissues.^39,65^ Consistent with this idea, we find that increased E-cadherin in E-cad OE embryos slows the speed of two steps during cell rearrangement: cell edge contraction and vertex resolution. In contrast, decreased E-cadherin in E-cad RNAi embryos also slows cell edge contraction but accelerates vertex resolution. Thus, vertex resolution rates show a monotonic decrease with E-cadherin levels, suggesting that the time to fully disassemble adhesions at a very short shrinking edge increases with the number of E-cadherin molecules at the contact. In contrast, cell edge contraction does not show a monotonic dependence on E-cadherin levels, indicating a more complex dependence of contraction rates on adhesion that might reflect multiple roles for E-cadherin in this process, e.g. in regulating the actomyosin contractile machinery, controlling adhesion strength, influencing junctional trafficking, and/or providing friction between cells. Future experimental studies of E-cadherin structure and function in cell rearrangement dynamics will be needed to explore how E-cadherin contributes to friction between cells.

Notably, while the ∼4-fold changes in E-cadherin levels in these studies tune aspects of cell packings and cell rearrangement dynamics, these changes have relatively weak effects on the overall morphogenetic process of body axis elongation. Although embryogenesis is disrupted in a large fraction of embryos as E-cadherin levels are significantly raised or lowered, if an embryo develops to the body axis elongation stage, it is very likely to elongate nearly fully and go on to hatch into a larva. These findings are consistent with the original characterization of the *shotgun* gene that encodes E-cadherin.^23,24^ *Shotgun* mutants displayed tissue cohesion defects specifically in regions of the embryo undergoing dynamic morphogenetic movements involving cell rearrangements,^24^ suggesting a minimum level of E-cadherin required for tissue cohesion. However, they also observed relatively uniform E-cadherin levels across the embryonic epithelium in wild-type embryos,^24^ suggesting that the precise level of E-cadherin in a region, so long as it is above the cohesion threshold, may not be specifically tuned for distinct morphogenetic processes. Our observations are also reminiscent of previous findings on the role of adhesion in cell sorting in the *Xenopus* embryo,^58^ indicating a minimum adhesion requirement for tissue cohesion but minimal effects on the dynamics of cell sorting. Taken together, our findings support the notion that there is a minimum threshold of E-cadherin levels required for tissue formation and integrity. Further, our results suggest that above this threshold, either overall body axis elongation is only weakly dependent on adhesion or that at a systems level the cellular contractile and adhesive machineries can adjust to a broad range of absolute E-cadherin levels to produce robust tissue elongation. The quantitative results in the present study will be useful for building and constraining models of epithelial mechanics that incorporate more complete pictures of cellular machineries and can recapitulate the remarkable robustness of morphogenetic processes.

## Materials and methods

Fly embryos were generated at 18°C or 23°C and analyzed at room temperature. E-cadherin levels were modulated by transgenic overexpression (BDSC #58445)^66,67^ or transgenic RNAi (BDSC #32904)^40^ using the Gal4/UAS system. E-cadherin levels were quantified in embryos fixed and stained using anti-DE-cadherin DCAD2 (DSHB). For live imaging, cell membranes were visualized with gap43:mCherry,^68^ E-cadherin with GFP-tagged DE-cadherin (BDSC #60584^69^ or BDSC #58445^66^), or myosin II with sqh:mCherry^60^. For fluorescence microscopy studies, embryos were imaged on a Zeiss LSM880 laser-scanning confocal microscope. Time-lapse movies were manually analyzed with ImageJ^70^ for quantifying E-cadherin levels; semi-automatically analyzed with SEGGA software in MATLAB^43^ for quantifying cell shapes, cell rearrangement rates, and cell areas; PIVlab Version 1.41 in MATLAB^71^ for quantifying tissue elongation; EPySeg^72^ or Cellpose 2.0^73^ for cell segmentations with a custom Python pipeline to quantify protein intensities; and custom code for quantifying the cell shape alignment index.^13,44^ Unless otherwise noted, error bars are the standard deviation (SD) and mean values from normal distributions were compared by one-way ANOVA and Tukey HSD tests. Statistical significance at the level of P<0.05 is denoted by a star.

## Acknowledgements

The authors thank the Bloomington *Drosophila* Stock Center (BDSC) for fly stocks; Mevludin Isic and Erik Boyle for assistance with data processing; Erika Kusaka for assistance with experiments. This work was supported by the National Institute of General Medical Sciences of the National Institutes of Health Award Number R35GM138380 to K.E.K. and Eunice Kennedy Shriver National Institute of Child Health & Development of the National Institutes of Health Grant F31HD105405 to C.C. K.E.K. holds a Burroughs Wellcome Fund Career Award at the Scientific Interface, NSF CAREER Award, Packard Fellowship, and Sloan Research Fellowship in Physics.

## Notes

### Competing Interest Statement

The authors have declared no competing interest.

